# Warm and cool temperatures decrease early-life telomere length in wild pied flycatchers

**DOI:** 10.1101/2025.05.02.651824

**Authors:** C Furic, C Marciau, B-Y Hsu, N Cossin-Sevrin, J Fleitz, S Reichert, S Ruuskanen, A Stier

## Abstract

Climate change represents a major challenge for avian species. It is characterized by an increase in average ambient temperatures, but also in occurrence of extreme weather events, such as heat weaves and cold snaps. These abrupt temperature changes can modify the immediate and long-term survival prospects of nestling birds, when their thermoregulatory capacities are still not fully developed. While immediate nestling survival can easily be measured, long-term survival is more challenging to evaluate. Early-life telomere length has been suggested as a potential biomarker of future fitness prospects. To evaluate the potential impact of changes in early-life temperature, we thus experimentally increased (*ca*. +2.8°C) and decreased (*ca*. −1.7°C) average nestbox temperatures in wild pied flycatchers (*Ficedula hypoleuca*) during nestling postnatal growth, and measured nestling telomere length before fledging. Shorter telomeres were observed in individuals exposed both to an experimental heating or cooling during growth. Our results suggest that long-term survival prospects or long-term performances of individuals exposed to abrupt changes in early-life temperature may be decreased.

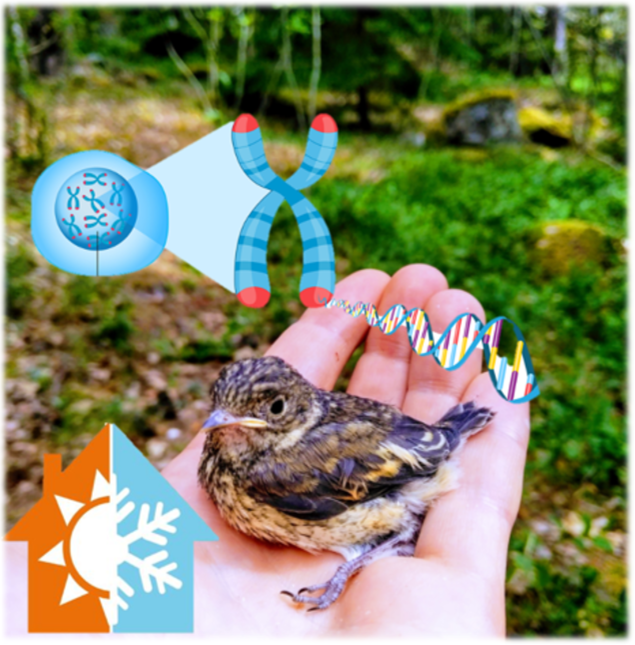

## Introduction

Ongoing climate change involves not only rising global mean temperatures but also an increase in the occurrence of extreme weather events such as cold snaps and heat waves (IPCC 2014). These abrupt temperature changes can cause direct mortality (McKechnie & Wolf 2009, McKechnie *et al*. 2021) but also sub-lethal consequences that may impact fitness because of delayed or long-lasting effects (Conradie *et al*. 2019). These extreme weather events can represent a major challenge for avian species, especially during early development when altricial nestlings are transitioning from an ectothermic stage to a fully endothermic stage prior to fledging (Price & Dzialowski 2018). As individuals do not fully regulate body temperature during these early-life stages (Boyles *et al*. 2011), their homeothermy depends on ambient temperature fluctuations and parental brooding throughout this thermoregulatory transition (Katzenberger *et al*. 2015). This temperature-sensitivity exposes individuals to short-term mortality risks caused by temperature variation (Hepp & Kennamer 2012, Moreno *et al*. 2015, Andreasson *et al*. 2018, Ospina *et al*. 2018) but also potential long-lasting or delayed effects on their health or reproduction (Metcalfe & Monaghan 2001, Preston *et al*. 2018) that may affect fitness (Greño *et al*. 2008, Hepp & Kennamer 2012, Berntsen & Bech 2016, Nord & Nilsson 2016, Andreasson *et al*. 2018). Previous studies have demonstrated the effect of heatwaves (Greño *et al*. 2008, Nord & Nilsson 2016, Andreasson *et al*. 2018) or cold snaps (Moreno *et al*. 2015, Ospina *et al*. 2018) events during pre- or postnatal stages on growth and parental brooding behavior (Rodríguez & Barba 2016). However, the potential long-term survival effects of such abrupt changes in temperature can be more difficult to characterize, because long-term recaptures rates are often very low for passerine birds (Andreasson *et al*. 2018, Stier *et al*. 2025). Heat waves or cold snap may have direct causal effects, but also indirect effects since changes in temperature are known to influence numerous biotic factors (*e*.*g*. food availability, parasite abundance; Schöll *et al*. 2016, Mennerat *et al*. 2021). Thus, testing for a causal effect of temperature requires experimental approaches, such as nest heating (*e*.*g*. Rodríguez & Barba 2016a, Andreasson *et al*. 2018) or cooling (*e*.*g*. Rodríguez & Barba 2016b, Corregidor-Castro *et al*. 2023).

To elucidate these potential long-term or delayed effects of early-life thermal environment, telomere length may be used as a molecular biomarker for future fitness prospects (Eastwood et al. 2019). Located at chromosome ends, telomeres serve as a protective role by preserving genome integrity and ensuring proper cellular division (Lange *et al*. 2006). It has been shown that telomeres progressively shorten with age (Remot *et al*. 2022), and that telomere length predicts survival prospects in some species at least (Wilbourn *et al*. 2018). Their rate of shortening can be accentuated by environmental challenges, particularly those encountered during early-life development, when telomere shortening is the most pronounced (Chatelain *et al*. 2019). For instance, telomere length may be affected by early-life temperature conditions, with experimental elevation of egg incubation temperatures leading to shorter telomeres (Vedder *et al*. 2018, Stier *et al*. 2020a, Hope *et al*. 2023). Yet, the effect of heatwaves and cold snaps on telomere length during postnatal stages have been less characterized. There is some evidence of adverse effects of natural or experimental hot conditions on telomere length in the wild (from postnatal d7 to d14: Stier *et al*. 2025; from postnatal d0 to d7: Eastwood *et al*. 2022), but no effect of experimental heatwaves on telomere length in captivity (from hatching to postnatal d12: Ton *et al*. 2023). To the best of our knowledge, the effect of postnatal coldsnap on telomere length has never been investigated, but two studies simulating parental neglects (*i*.*e*. cooling down nestling for up to 1 hour per day for *ca*. the first week post-hatching) found no significant effect on telomere length (Lynn *et al*. 2022, Ghimire *et al*. 2025). Accelerated telomere attrition can be linked to fast growth and rapid cellular proliferation (Monaghan & Ozanne 2018), oxidative damage (Reichert & Stier 2017; Armstrong & Boonekamp 2023), or disruptions in metabolic balance (Casagrande & Hau 2019). All these mechanisms may underlie potential effects of early-life temperature on telomere length.

In this study, we investigated the effects of experimental increase and decrease in nest temperature during early postnatal development on telomere length in wild pied flycatchers (*Ficedula hypoleuca*). As study years also differed in terms of ambient thermal conditions (see results), we also used this opportunity to investigate the potential relationship between yearly ambient temperature and telomere length. Age- and stress-related telomere shortening have been previously demonstrated in this species and its close relative, the collared flycatcher (*Ficedula albicollis)* (Kärkkäinen *et al*. 2019, 2020, 2021, 2022, Stier *et al*. 2020b). Investigating experimentally the effects of heatwave and coldsnap on postnatal telomere length in a free-living species should help to determine the potential long-term or delayed effects of extreme weather events on fitness. We predict that being exposed to early-life hot environmental conditions should lead to shorter telomeres, as found in Stier *et al*. (2025) and Eastwood *et al*. (2022). However, experiencing a coldsnap may lead either to shorter telomeres, if the coldsnap triggers an increase in metabolic rate or energetic imbalance (Casagrande & Hau 2019, Casagrande *et al*. 2020), or to longer telomeres, if the coldsnap reduces the metabolic rate of partly ectothermic nestlings (Friesen *et al*. 2022).

## Material and methods

### General field procedures

Two nest temperature manipulations were conducted in a nestbox population of pied flycatchers located in the island of Ruissalo (Turku, Finland, location: 60°25’N, 22°10’E). In 2018, a heating experiment was performed, (data partly published in Hsu *et al*. (2020)), and in 2019 a cooling experiment was performed. Nestbox temperature was recorded every 3 minutes with an Ibutton thermochron (0.0625°C accuracy) installed *ca*. 5 cm above the rim of the nest. On the second day after hatching (d2), each individual was identified by clipping nails and half of the nestlings (minimum 2 individuals) with the same age and weight were swapped with another nest for other experimental purposes (*i*.*e*. cross-fostering to disentangle environmental and genetic effects).

### Heating experiment (2018)

In 2018, nestbox temperature was increased for half of the boxes (hereafter, ‘heated nests’) between postnatal d2 and d8. Heating pads (UniHeat 72h, USA) were installed under the ceiling and replaced every 2^nd^ day in the morning. The other half was considered as controls (hereafter, ‘non-heated nests’), non-functional heating pads were used and nests were visited every 2^nd^ day as well to standardize disturbance. At d8 after hatching, individuals were identified and ringed, and at postnatal d13 one blood sample of *ca*. 40μl was collected from the brachial vein using heparinized capillaries. A total of 97 nestlings from 41 nests were analyzed, specifically 42 individuals from 21 heated nests and 55 individuals from 20 non-heated ones. When possible, the nestlings selected for telomere analysis included (1) both a female and male nestling from each nest (2) one of genetic origin and one foster nestling, and otherwise were randomly selected. Since females can actively brood nestlings until postnatal d6 or d7 (Lundberg & Alatalo 1992), they could potentially change their behavior depending on the heating treatment, but no evidence was found in this experiment (previously published in Hsu *et al*. (2020)).

### Cooling experiment (2019)

In 2019, nestbox temperature was decreased for half of the boxes (hereafter, ‘cooled nests’) between d7 and d13 after hatching of the nestlings. The timing of the experiment was chosen to avoid any potential confounding effects of brooding behavior by the female, and knowing that pied flycatchers are not yet fully capable of thermoregulation at d7 (Lundberg & Alatalo 1992). Two frozen (−20°C) cooling pads (VWR 26.7*14*3.2*cm, 909g) were attached on each outer side of the nestbox (protected on the outside with insulating material) and replaced every morning. The other half (hereafter, ‘non-cooled nests’) was equipped with the same pads, but unfrozen, and visited every day too. Before the treatment, at d7 after hatching, individuals were identified and ringed. Morphological measurements (body mass and wing length) were conducted at postnatal d13, and one blood sample of *ca*. 40μl was collected from the brachial vein using heparinized capillaries. A total of 56 nestlings from 48 nests were analyzed, including 25 individuals from 23 cooled nests and 31 individuals from 25 non-cooled nests. The nestlings were selected for telomere analysis by trying to balance the number of female and male nestlings, and having one of genetic origin and one foster nestling.

### Relative telomere length analysis

Blood samples were stored in a cooler (*ca*. 4°C) during the day of collection in the field, and were transferred to a −80°C freezer at the end of the day. We measured relative telomere length using a real-time quantitative PCR (qPCR) assay previously used and validated in pied flycatcher using the TRF method (Kärkkäinen *et al*. 2019, 2021). DNA was extracted from whole blood samples using the standard salt extraction alcohol precipitation method (Aljanabi & Martinez 1997). DNA concentration and purity were assessed spectrophotometrically (ND-1000-Spectrophotometer, Thermo Fisher Scientific, Waltham, MA, USA), and the DNA extract was considered good if purity (260/280 > 1.8 and 260/230 > 2.0) and quantity ([DNA] > 40 ng.μL^-1^) criteria were met. DNA integrity was checked using gel electrophoresis (50 ng of DNA, 0.8% agarose gel at 100 mV for 60 min, Midori Green staining (NIPPON Genetics Europe, Düren, Germany). Each sample was then diluted to 1.2 ng.μl^-1^, and the final reaction contained 6 ng of DNA, 300nM of forward and reverse primers and SensiFAST SYBR Lo-ROX master mix (Bioline, London, UK) for a total volume of 12 μl. Tel1b was used as a forward telomere primer (5′-CGGTTTGTTTGGGTTTGGGTTTGGGTTTGGGTTTGGGTT-3′), Tel2b as the reverse one (5′-GGCTTGCCTTACCCTTACCCTTACCCTTACCCTTACCCT-3′) and RAG1 as a single copy gene (forward primer 5′-GCAGATGAACTGGAGGCTATAA-3′ and reverse primer 5′-CAGCTGAGAAACGTGTTGATTC-3′) following Kärkkäinen *et al*. (2021). The qPCR conditions started with an initial denaturation (3 min at 95°C), then 40 cycles with first step of 10s at 95°C, second step of 15s at 58°C and third step of 10s at 72°C, with melting curve analysis in the end performed by QuantStudio™ 12K Flex Real-Time PCR System (Thermo Fisher Scientific, Waltham, MA, USA) using 384-well plates. Each plate contained negative control and three internal standards. Samples from the different experimental treatments were distributed evenly across all plates. To determine the baseline fluorescence, the quantification cycle values and the qPCR efficiencies, we used LinRegPCR (Ruijter *et al*. 2009). Relative telomere length was calculated based on plate-specific efficiencies (average ± SE and [range] for telomere and RAG1 reactions were, respectively 95.1 ± 1.6% [90.1 – 103.3%] and 101.7 ± 1.8% [94.4 – 105.8%]) using the mathematical model presented in Pfaffl (2001). Technical repeatability based on triplicate measurements of telomere length was 0.74 (95% Cl [0.69, 0.79], *p* < 0.001), and inter-plate repeatability (calculated by using the average of triplicates) based on 37 samples measured on more than one plate was 0.80 (95% Cl [0.67, 0.93], *p* < 0.001). Samples have been analyzed over 16 qPCR plates in total. Molecular sexing was conducted using qPCR (adapted from Ellegren & Fridolfsson 1997, Chang *et al*. 2008; see details in Hsu *et al*. (2020)).

### Statistical analysis

Statistical analysis was carried out using R software version 4.3.2 (R Core Team 2014). Normality and homogeneity of residual variances were checked using the R package DHARMa (Hartig 2022).

#### Effect of the experimental manipulations and year on nestbox temperature

For each year, to evaluate nestbox temperature (*i*.*e*. hourly average) differences among groups, we used linear mixed-effects models (LMMs - implemented in the lme4 package (Bates *et al*. 2015), Satterthwaite’s degrees of freedom), with group (2018: heated vs. non-heated; 2019; cooled vs. non-cooled) and time of day (*i*.*e*. treated as a categorical variable) treated as fixed effects. We included random intercepts for date (to account for temporal autocorrelation) and rearing nestbox (to control for repeated measurements).

To investigate natural variation in ambient and nestbox temperatures between 2018 and 2019 (for each year, between June 4th to July 12th), LMMs were used on hourly air temperature (provided by the Finnish Meteorological Institute and collected by Artukainen weather station in Turku, 2 to 4 km from the study sites) and nestbox temperature (*i*.*e*. only non-heated and non-cooled), with year and time of day included as fixed effects, and date modeled as a random effect to account for repeated measurements within days. Model parameters were estimated using restricted maximum likelihood (REML).

#### Effect of the experimental manipulations of temperature on body mass

The effect of heating treatment on growth was analysed previously and no effect was detected at any age for body mass (Hsu *et al*. 2020). We conducted similar analyses for the cooling experiment, and no significant effects of the treatment were detected for body mass (*t* = 0.39, *P* = 0.70).

#### Effect of the experimental manipulations of temperature on telomere length

A LMM on relative telomere length for each dataset was conducted separately to assess the effects of the treatment (heated vs. non-heated nests; cooled vs. non-cooled nests) and other traits of interest (*i*.*e* sex, cross fostering, hatching date, brood size and body mass). Nest of origin, nest of rearing and the qPCR plate identities were integrated in the model as random intercepts. No significant sex-dependent effects of temperature treatments on telomere length were found (*P* > 0.12), and thus interactions between sex and treatment were removed from final models to enable proper interpretation of the main effects.

#### Relationships between nestbox temperature and telomere length

Although analyzing experimental data with the above-mentioned treatment *vs*. control comparisons is the most appropriate way to test for a causal effect of temperature on telomere length, we also ran correlative analyses on both experiments, replacing experimental treatment by the temperature measured inside each nestbox. We ran 3 models per experiment, one for mean temperature, one for minimum daily temperature (averaged over the 7 days of treatment), and one for maximum daily temperature (averaged over the 7 days of treatment). These models had the exact same structure as the ones described above, except for the replacement of treatment by nestbox temperature (mean, minimum or maximum).

#### Year effect on telomere length

As experimental manipulations differed markedly between years (*e*.*g*. timing of temperature manipulation), we cannot directly compare the two experiments. Yet, environmental conditions (including ambient temperature, see results) differed between the two study years, and thus comparing telomere length of control individuals between years (*i*.*e*. non-heated from 2018 and non-cooled from 2019) may provide information regarding the response to longer-term natural variation in temperature. This was conducted using a linear mixed model relatively similar to the ones described above, with telomere length as a response variable, the different traits of interest (*i*.*e* year, sex, cross fostering, hatching date, brood size and body mass) as fixed effects and three random intercepts (*i*.*e* nest of origin, nest of rearing and qPCR plate identities). Only 2 adults bred both years, which is insufficient to include the identity of the parents as a random intercept.

## Results

### Nestbox temperature: experimental effects and natural variation between years

In 2018, the temperature in heated nest boxes was on average higher than in non-heated ones (estimate: +2.83°C, 95% CI: [1.73; 3.53], t = 6.09, p < 0.001, Fig 1A). In 2019, the temperature in cooled nestboxes was on average lower than in non-cooled nestboxes (estimate: −1.72°C, 95% CI: [-1.56; −0.45], t = −3.52, p < 0.01, Fig 1 B). Yet, there were in both cases strong interactions between time of day and experimental treatments (2018: F_23.1_=31.2, p<0.001; 2019: F_23.1_=103.7, p<0.001), with treatment effect being more pronounced during mid-day (Fig. 1A and 1B). For instance, nestbox temperature at 16:00 was *ca*. 3.0°C higher in heated nests (Fig. 1A) and *ca*. 2.0°C lower in cooled nests (Fig. 1B).

**Fig 1:**
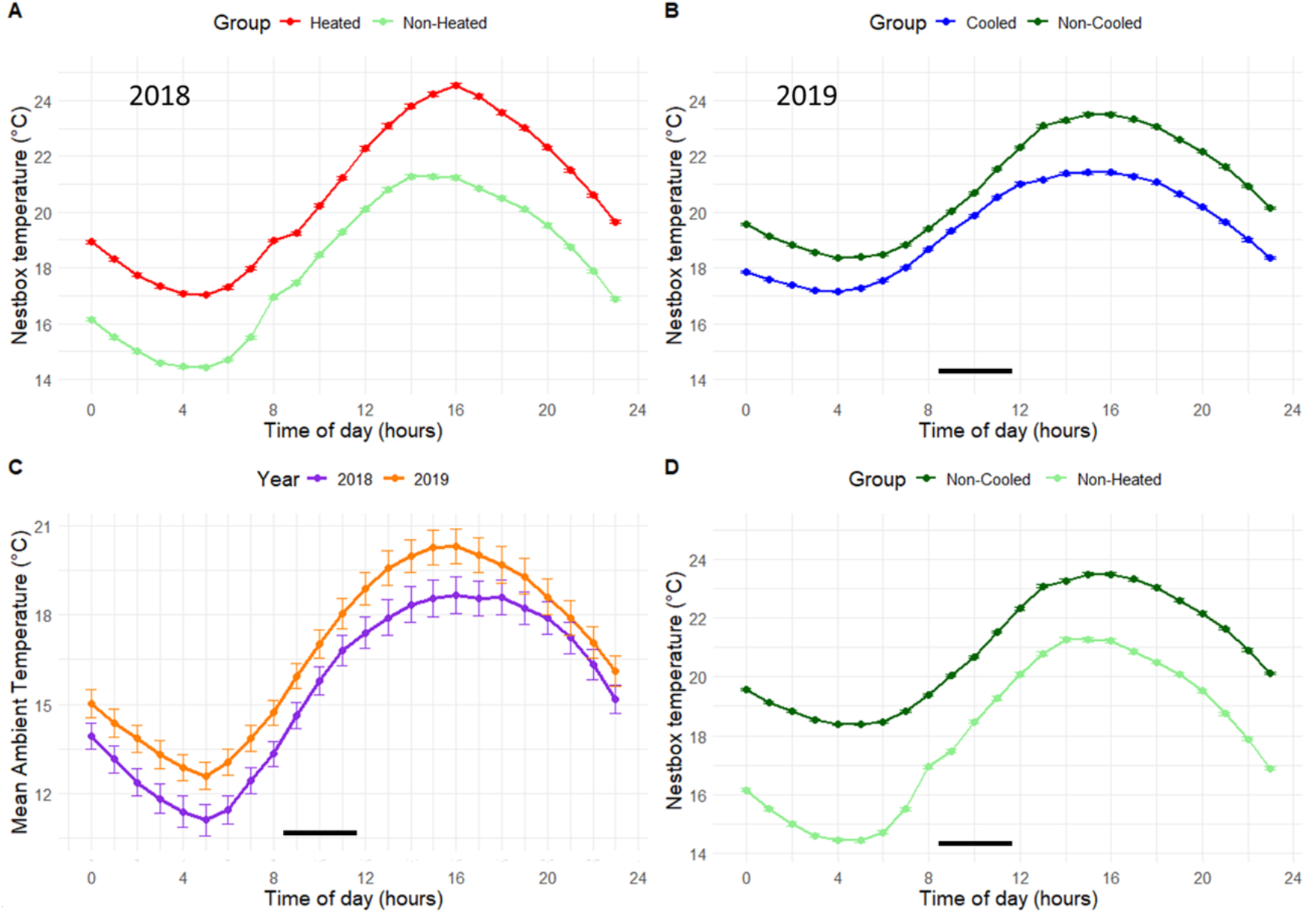
Experimental and natural variation in nestbox and ambient temperatures: effect of experimental heating (A) and cooling (B) on nestbox temperature; natural variation between years in ambient (C) and nestbox (D) temperatures. The black bar represents the time of cooling or heating pads replacement. Hourly mean ± SE are reported for 41 nests in 2018 (21 heated vs; 20 non-heated) and 48 nests in 2019 (23 cooled vs. 25 non-cooled).

During the experiments (June-July), ambient temperature was significantly higher in 2019 than 2018 (estimate: +1.54°C, 95% CI: [0.93; 2.15], t = 4.98, p < 0.001, Fig 1 C). This temperature difference was also observed in nestbox temperatures between non-heated and non-cooled groups with higher temperature in 2019 than 2018 (estimate: 2.16°C, 95% CI: [1.08; 3.24], t = 3.92, p < 0.01, Fig 1 D).

### Effect of experimental manipulations and natural variation in nestbox temperature on telomere length

Nestlings from heated nests had significantly shorter telomeres than non-heated ones (LMM: *t*_*26*.*39*_ = 2.85, *P* = 0.008; Fig.2A; Table 1A) and nestlings from cooled nests had shorter telomeres compared to the ones from non-cooled nests (LMM: *t*_*35*.*26*_ = 2.23, *P* = 0.032; Fig. 2B; Table 1B). Replacing heating treatment by the actual nestbox temperature revealed no significant relationship between telomere length and mean (*β* ± SE = −0.012 ± 0.015, *P* = 0.43, Table S1A) or minimum temperature (*β* ± SE = −0.005 ± 0.015, *P* = 0.72, Table S2A), but there was a non-significant trend for higher maximum temperature to be associated with shorter telomeres (*β* ± SE = −0.017 ± 0.009, *P* = 0.073, Table S3A). Replacing cooling treatment by the actual nestbox temperature revealed that cooler mean temperature was associated with shorter telomeres (*β* ± SE = 0.068 ± 0.032, *P* = 0.043, Table S1B), but no significant relationship was found with minimum (*β* ± SE = 0.046 ± 0.032, *P* = 0.17, Table S2B) nor maximum temperature (*β* ± SE = 0.016 ± 0.017, *P* = 0.36, Table S3B).

**Table 1:** Summary of the linear mixed models testing the effects of experimental temperature manipulation ((A) heating treatment in 2018, (B) cooling treatment in 2019) and relevant covariates on relative telomere length of pied flycatcher nestlings at day 13 after hatching. Non-significant interactions between treatment and sex were removed from final models, but are presented in the table for completeness. Singular fit partly occurred for the nest of rearing and the identity of qPCR plate for the heating experiment, and nest of origin for the cooling experiment, but they were kept as random intercepts as they were part of the experimental design.

**(A) Experimental heating (2018, N = 97 nestlings)**
| Random effects: |  | Variance |  |  |  |
| --- | --- | --- | --- | --- | --- |
| Nest of origin<br>(N=26) | Intercept | 0.0028 |  |  |  |
| Nest of rearing<br>(N=41) | Intercept | 0 |  |  |  |
| Plate qPCR (N=6) | Intercept | 0 |  |  |  |
| Residual |  | 0.0474 |  |  |  |
| Fixed effects: | Estimate | Std. Error | t | df | P (>\|t\|) |
| Intercept | 1.120 | 0.346 | 3.24 | 63.65 | 0.002 |
| Heating (Non-heated) | 0.141 | 0.049 | 2.85 | 26.39 | 0.008 |
| Sex (Male) | -0.054 | 0.046 | -1.18 | 88.13 | 0.24 |
| Cross-fostering<br>(Yes) | 0.054 | 0.050 | 1.08 | 59.94 | 0.29 |
| Hatching date | 0.002 | 0.004 | 0.62 | 46.89 | 0.54 |
| Brood size | 0.013 | 0.023 | 0.57 | 39.26 | 0.57 |
| Body mass | -0.018 | 0.018 | -0.99 | 66.03 | 0.33 |
| ..... |  |  |  |  |  |
| Heating (Non-heated) × Sex<br>(Male) | -0.145 | 0.093 | -1.54 | 88.25 | 0.12 |
| ----- |  |  |  |  |  |
| Sample size: N = 55 nestlings from non-heated nest, N = 42 nestlings from heated nest |  |  |  |  |  |

**(B) Experimental cooling (2019, N = 56 nestlings)**
| Random effects: |  | Variance |  |  |  |
| --- | --- | --- | --- | --- | --- |
| Nest of origin<br>(N=29) | Intercept | 0 |  |  |  |
| Nest of rearing<br>(N=48) | Intercept | 0.005 |  |  |  |
| Plate qPCR (N=10) | Intercept | 0.019 |  |  |  |
| Residual |  | 0.051 |  |  |  |
| Fixed effects: | Estimate | Std. Error | t | df | P (>\|t\|) |
| Intercept | 1.078 | 0.572 | 1.89 | 39.53 | 0.069 |
| **Cooling (Non-cooled)** | **0.156** | **0.070** | **2.23** | **35.26** | **0.032** |
| Sex (Male) | 0.020 | 0.072 | 0.27 | 44.48 | 0.79 |
| Cross-fostering (Yes) | -0.019 | 0.070 | -0.27 | 46.68 | 0.79 |
| Hatching date | 0.013 | 0.008 | 1.61 | 14.77 | 0.129 |
| Brood size | 0.033 | 0.030 | 1.11 | 30.21 | 0.28 |
| Body mass | -0.035 | 0.031 | -1.11 | 42.97 | 0.27 |
| Cooling (Non-cooled) × Sex (Male) | -0.034 | 0.148 | -0.23 | 44.98 | 0.82 |
| Sample size: N = 31 nestlings from non-cooled nest, N = 25 nestlings from cooled nest |  |  |  |  |  |

**Fig. 2:**
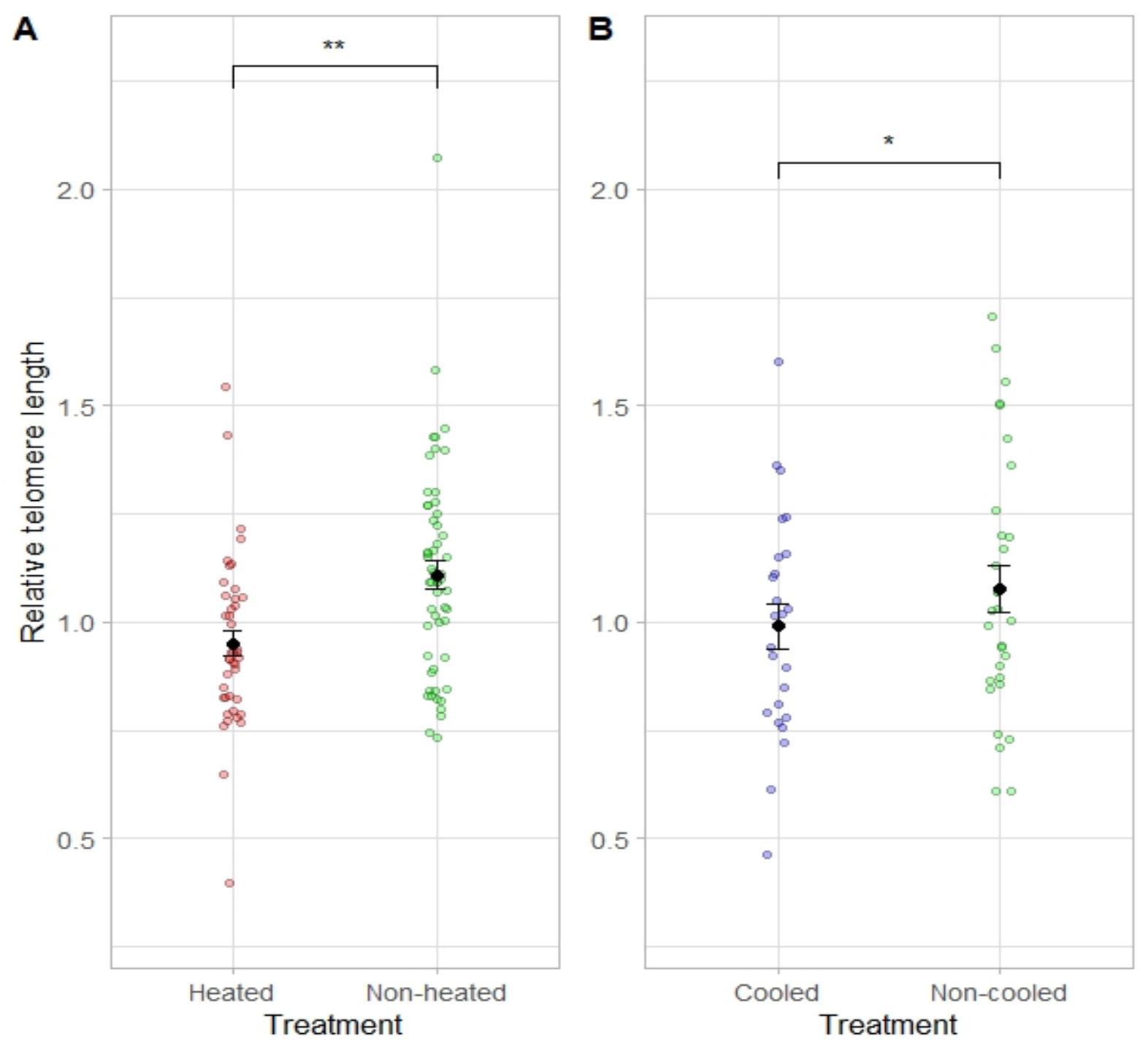
Effect of experimental heating (A) and cooling (B) on relative telomere length in pied flycatcher nestlings at day 13 after hatching. Individual raw data points are presented along with estimated marginal means ± SE from statistical models described in Table 1.

### Natural variation in telomere length and body mass between years

We observed no significant association between year and telomere length for control individuals (LMM: *t*_*5*.*00*_ = −0.31, *P* = 0.77; Fig. 3, Table 2), as well as no significant year effect on nestling body mass (LMM: *t* = 0.30, *P* = 0.76).

**Table 2:**
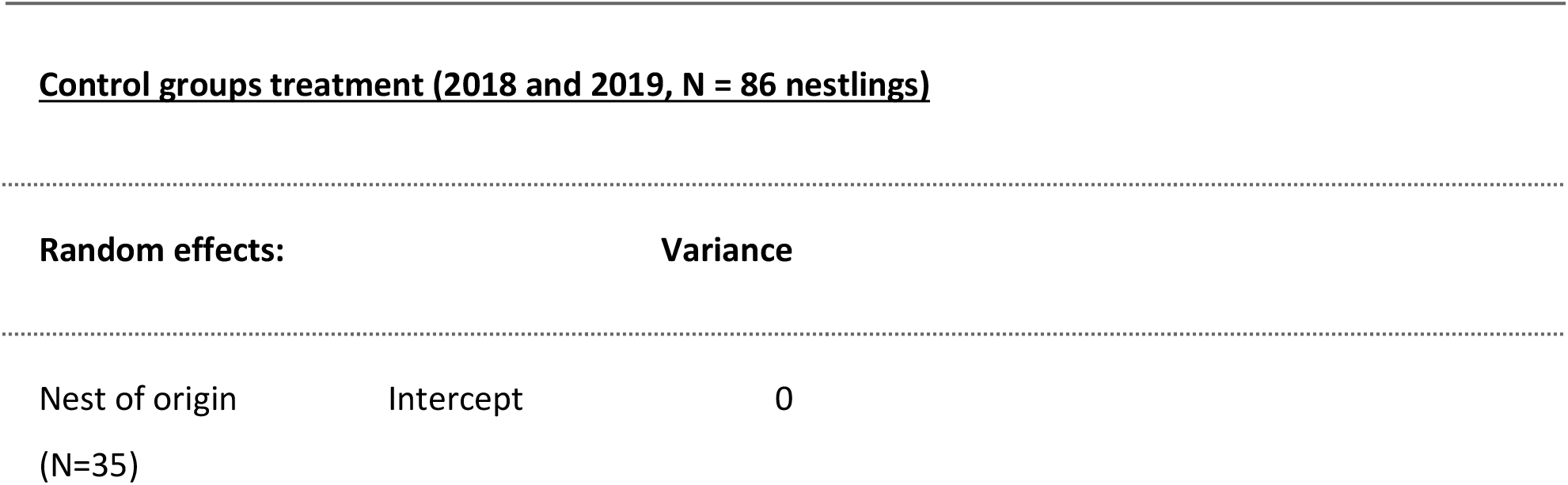
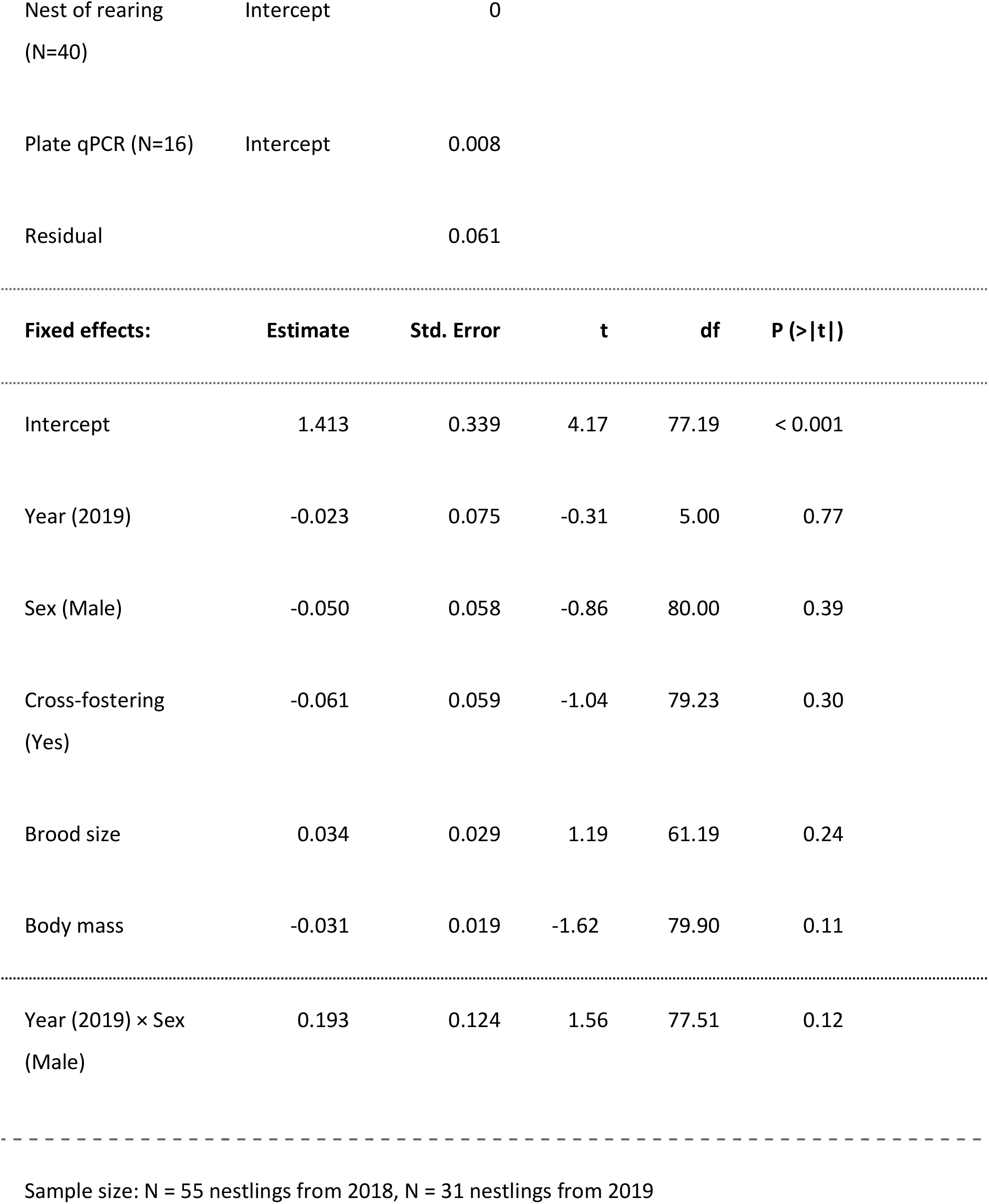
Summary of the linear mixed model testing the effects of the year (2018 vs. 2019) and relevant covariates on relative telomere length of pied flycatcher nestlings at day 13 after hatching from control groups (non-heated from 2018 and non-cooled from 2019). Non-significant interaction between year and sex was removed from the final model, but is presented in the table for completeness. Boundary singular fit occurred for the nest of rearing and nest of origin, but they were kept as a random intercept since it was part of the experimental design.

**Fig. 3:**
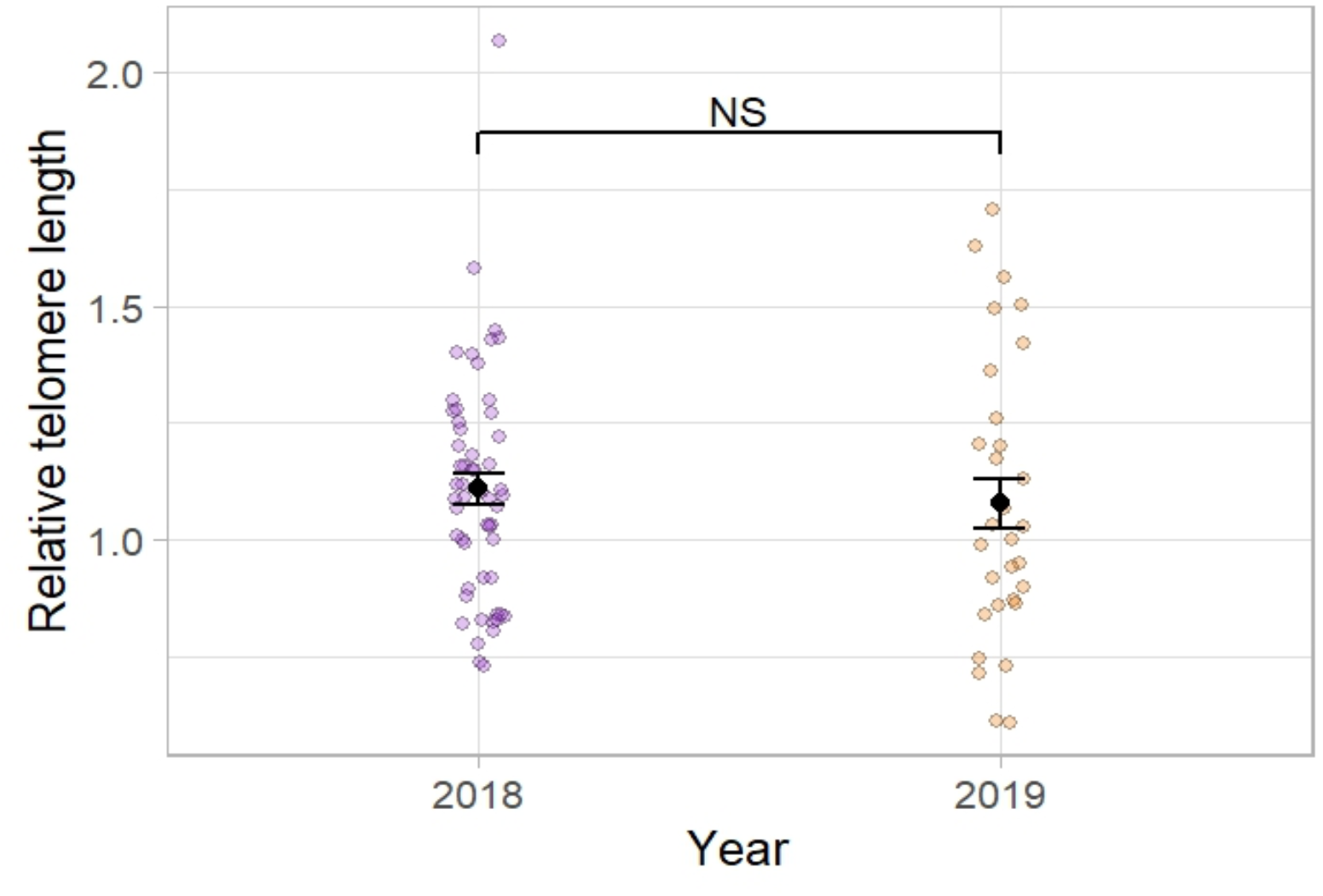
Effect of year on relative telomere length in pied flycatcher nestlings at day 13 after hatching from control nests. Individual raw data points are presented along with estimated marginal means ± SE from statistical models described in Table 2.

## Discussion

In this study, we demonstrated that exposure to experimental changes in environmental temperature (warmer or cooler) during early-life can significantly shorten telomeres in pied flycatcher nestlings. This suggests that the relationship between ambient temperature and telomere length in correlative analyses could be expected to be non-linear, following an inverted U-shape, as already suggested in ectotherms (Axelsson *et al*. 2020, Friesen et *al*. 2022) and to a lesser extent in birds (Pepke *et al*. 2022). Since avian altricial nestlings transition from ectothermy to endothermy during development (Price & Dzialowski 2018), these results echo the effects of experimental increase of temperature on telomere lengths in ectotherms, such as fishes (Siberian sturgeon (Simide *et al*. 2016), Atlantic Salmon at the embryo stage (McLennan *et al*. 2018)) or squamates (Desert Agama (Zhang *et al*. 2018)). To the best of our knowledge, only one cooling experiment has been conducted in ectotherms, showing no significant effect of a transition from 30°C to 20°C (Eastern mosquitofish (Rollings *et al*. 2014). Yet, it has been recently shown in amphibians that cooler temperature, for a similar growth rate, is associated with shorter telomeres (Burraco *et al*. 2023). So, our observations of shorter telomeres in response to experimental heating and cooling seem consistent with an ectothermic-like phenotype. Indeed, high temperature has also been shown to accelerate telomere shortening in ectothermic or partly ectothermic avian embryo and nestlings (Stier *et al*. 2020a, Stier *et al*. 2025, Eastwood *et al*. 2022), but also in endotherms such as humans (Ni *et al*. 2022) and domestic chicken (Musa *et al*. 2025). Future studies investigating temperature effects in fully endothermic avian models (*e*.*g*. after fledging) are needed to establish the potential role of (partial) ectothermy on the effects of temperature on telomere shortening.

In contrast with the results stemming from our experimental manipulation, we observed that while the mean ambient temperature of 2019 was 1.5°C higher than in 2018 (and nestbox temperature 2.2°C higher), there was no significant year effect on telomere length when comparing the data from control broods. This suggests that long-term variations in temperature may have less effect on telomere length than short-term changes mimicked by our experimental conditions, although this remains to be experimentally demonstrated. Indeed, an alternative explanation could be that the higher temperature in 2019 contributed to accelerate telomere shortening, but that higher food availability, usually resulting from warmer environmental temperatures (Avery & Krebs 1984, Veistola *et al*. 1997), compensated for any temperature-related shortening. It has for instance been shown that better nutritional conditions can limit telomere loss in great tit nestlings (Casagrande *et al*. 2023).

The temperature-related telomere shortening we observed in this study in response to experimentally increased or decreased temperature may be explained by several potential mechanisms. First, telomere shortening has been shown to be accentuated by fast growth and rapid cellular proliferation (Monaghan & Ozanne 2018). This seems unlikely to explain the effects we observed on telomeres since experimental heating (Hsu *et al*. 2020) or cooling (present study) had no detectable effects on body mass. Second, telomere shortening can be accelerated by increased oxidative stress levels (Reichert & Stier 2017), and acute heat or cold exposure have been shown to potentially increase oxidative stress levels (Costantini *et al*. 2012, Stier *et al*. 2014). Yet, this seems unlikely to explain the effects we observed on telomeres since experimental heating had no detectable effects on oxidative stress levels (Hsu *et al*. 2020). Third, telomere shortening can be accelerated by energetic imbalance, because of potential trade-offs between immediate and long-term survival during challenging environmental conditions (Casagrande & Hau 2019). This has been suggested to be mediated by glucocorticoid hormone (*i*.*e*. “stress hormone”) effects on cellular signaling via the mTOR pathway (Casagrande & Hau 2019). Since abrupt changes in temperature can increase glucocorticoid levels in altricial nestlings (Jessop *et al*. 2016, De Bruijn & Romero 2018), it is possible that the telomere shortening could be explained by this mechanism, as already suggested in an experimental heating study in great tit (Stier *et al*. 2025). Changes in ambient temperature can potentially influence body temperature, as for instance found in an experimental heating study in blue tits (Andreasson *et al*. 2018), with such effect being of higher magnitude at earlier (*i*.*e*. more ectothermic) postnatal stages. Changes in body temperature could be linked to energetic imbalance and glucocorticoid secretion in explaining temperature-dependent telomere shortening. Hypothetically, changes in body temperature could also be linked to telomere attrition via effects in telomerase (enzyme responsible for telomere elongation), since telomerase activity has been suggested to be temperature-sensitive (Friesen *et al*. 2022). Measuring glucocorticoid levels, mTOR signaling, body temperature and telomerase activity would be required to test these hypotheses.

To conclude, we have demonstrated that experimental increase or decrease in early-life temperature can shorten telomeres, which may impact long-term survival and fitness. Yet, directly measuring effects of experimental temperature manipulations on fitness would be needed to confirm this hypothesis, which is challenging in small passerine species having very low survival rates after fledging. Consequently, investigating those effects in long-lived species with high recruitment rate and known relationship between telomere length and fitness related traits (*e*.*g*. Alpine swift, Bize *et al*. 2009) would seem promising to better understand the effects of heatwaves and cold-snaps during early-life on telomeres and fitness.

## Author contribution

SRu, AS, CM and BYH designed the studies and conducted fieldwork. CM, AS and NCS conducted laboratory work. CF, JF and AS analyzed the data. CF, JF & AS wrote the manuscript, with input from SRu, SRe, NCS & CM. SRu and SRe acquired funding. SRu, SRe and AS supervised early-career researchers involved in this study.

## Acknowledgements

We are grateful to Lucas Bousseau, Thomas Rossille, Hsiao-Yin Liu, Päivi Kotitalo, Mélanie Crombecque, Axelle Delaunay, Lotta Holmen and Mikaela Hukkanen for their help in the field. The laboratory facilities were supported by Biocenter Finland.

## Data accessibility

All data are accessible at https://figshare.com/s/eb091d4259185e8646a6 (doi: 10.6084/m9.figshare.28844615)

## Ethical permits

All procedures were approved by the Animal Experiment Committee of the State Provincial Office of Southern Finland (license numbers ESAVI/2902/2018 & ESAVI/5718/2019) and by the Environmental Center of Southwestern Finland (license numbers VARELY/549/2018 & VARELY/924/2019) granted to SRu.

## Significance statement

Rapid environmental change is exposing wildlife to increasingly frequent and intense weather extremes, including heatwaves and cold snaps, which are particularly likely to impact sensitive early-life stages. Our results in pied flycatcher nestlings demonstrate that both experimentally elevated and reduced nest temperature during early life significantly shorten telomeres (i.e. a biomarker of long-term survival and fitness prospects). This study offers novel and ecologically meaningful insights into how environmental stress linked to climate change may shape avian life-history trajectories.

---

**Supplementary Table S1:**
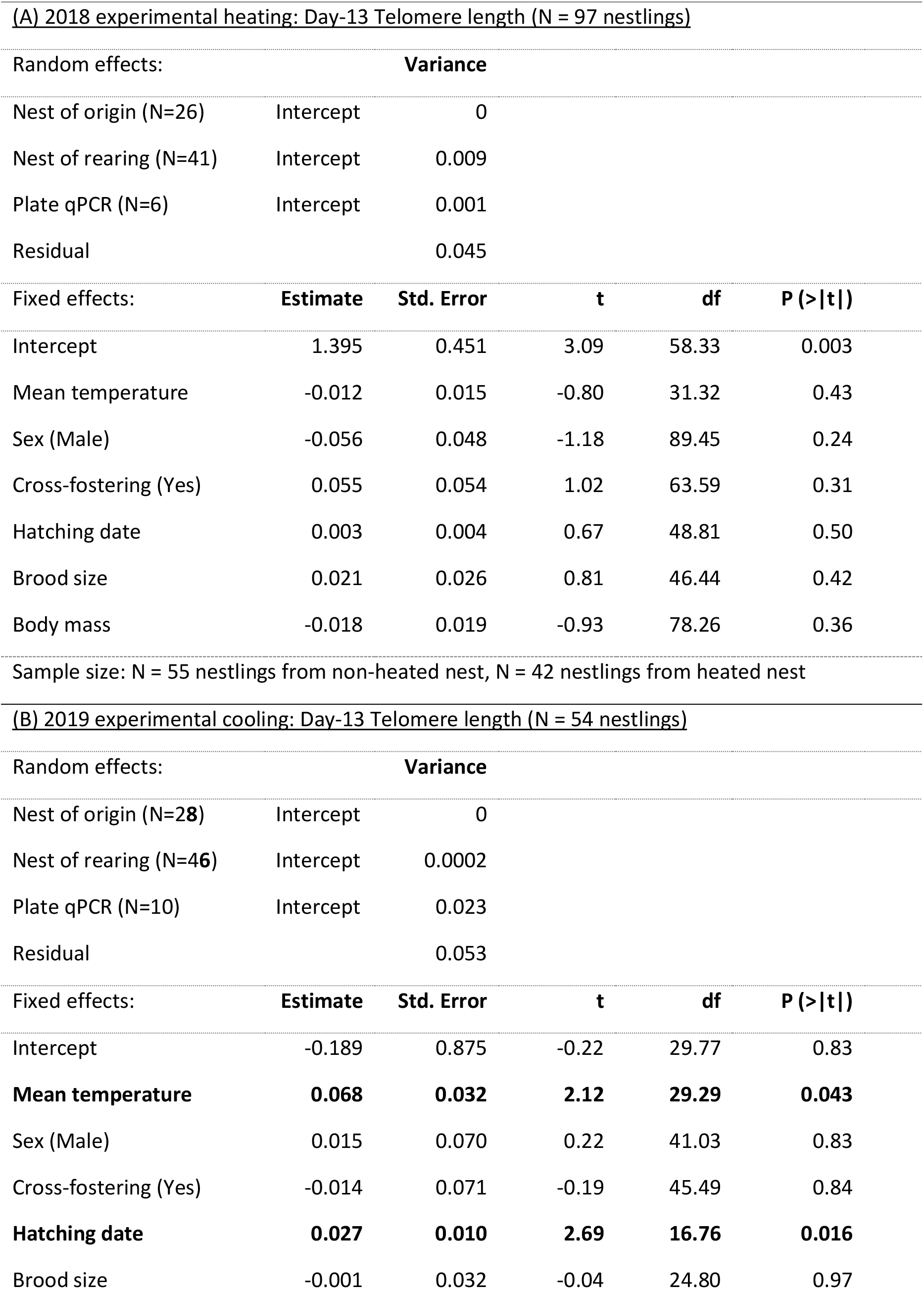
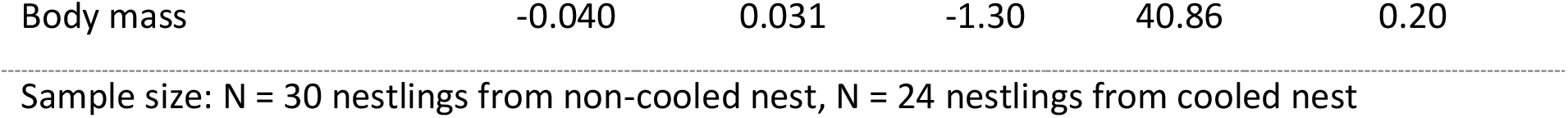
Summary of the linear mixed models testing the relationship between telomere length of nestlings at day 13 after hatching and <u>mean</u> nestbox temperature. ((A) 2018 heating experiment, (B) 2019 cooling experiment) while controlling for other variables.

**Supplementary Table S2:**
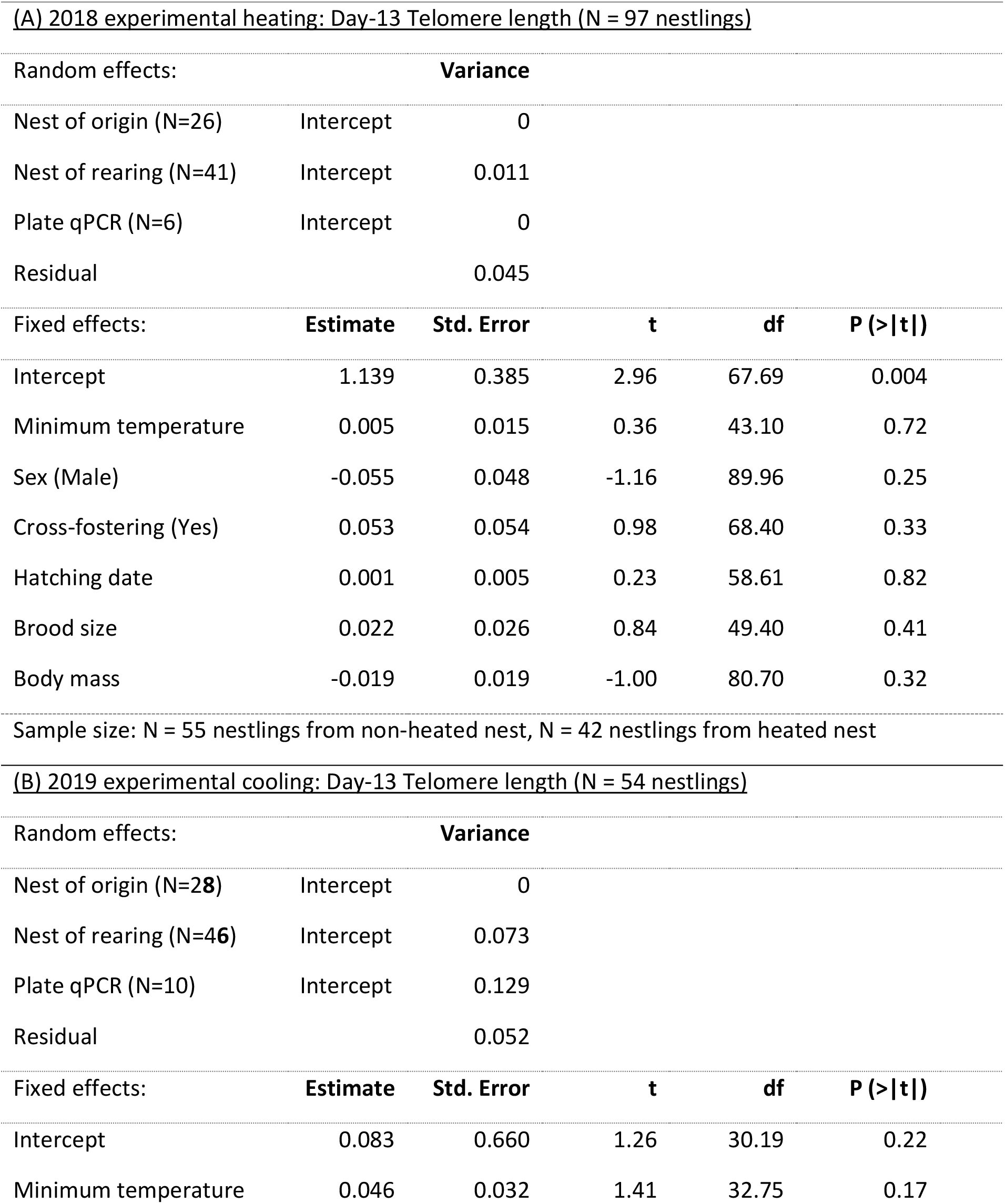
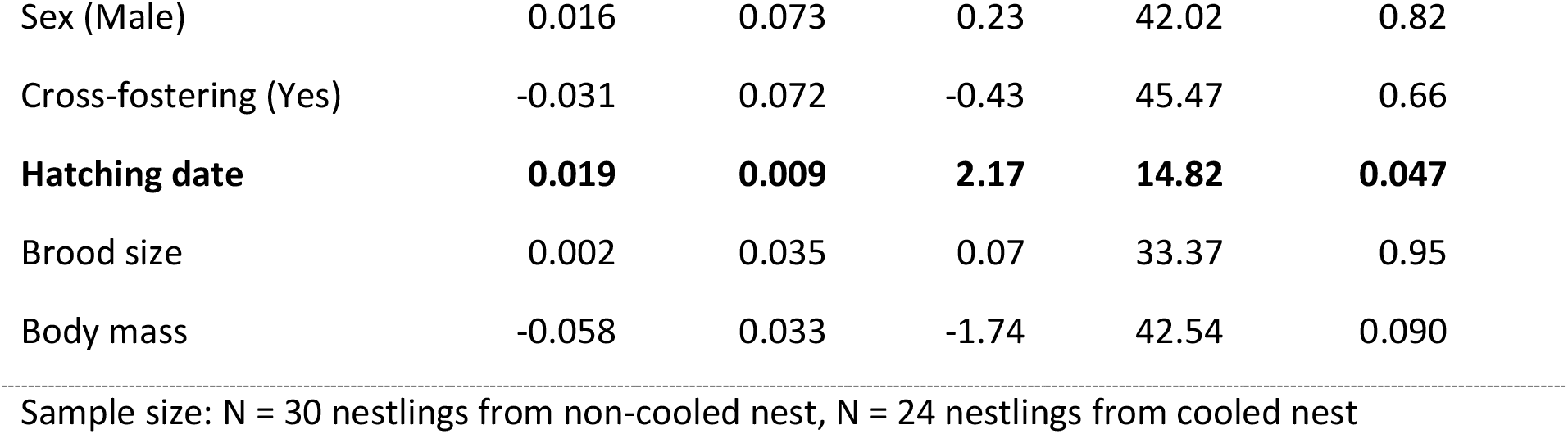
Summary of the linear mixed models testing the relationship between telomere length of nestlings at day 13 after hatching and <u>minimum</u> nestbox temperature. ((A) 2018 heating experiment, (B) 2019 cooling experiment) while controlling for other variables.

**Supplementary Table S3:**
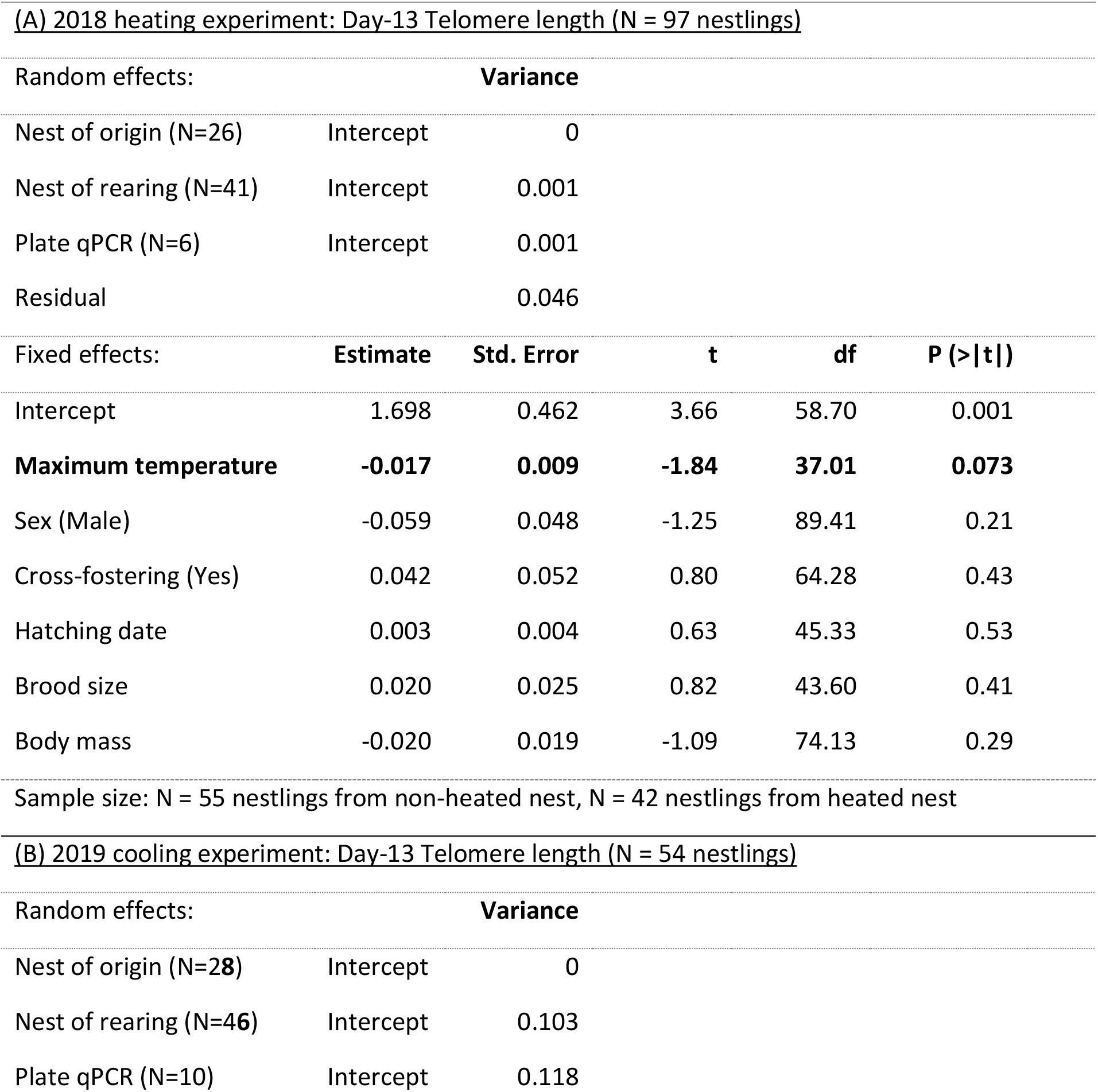
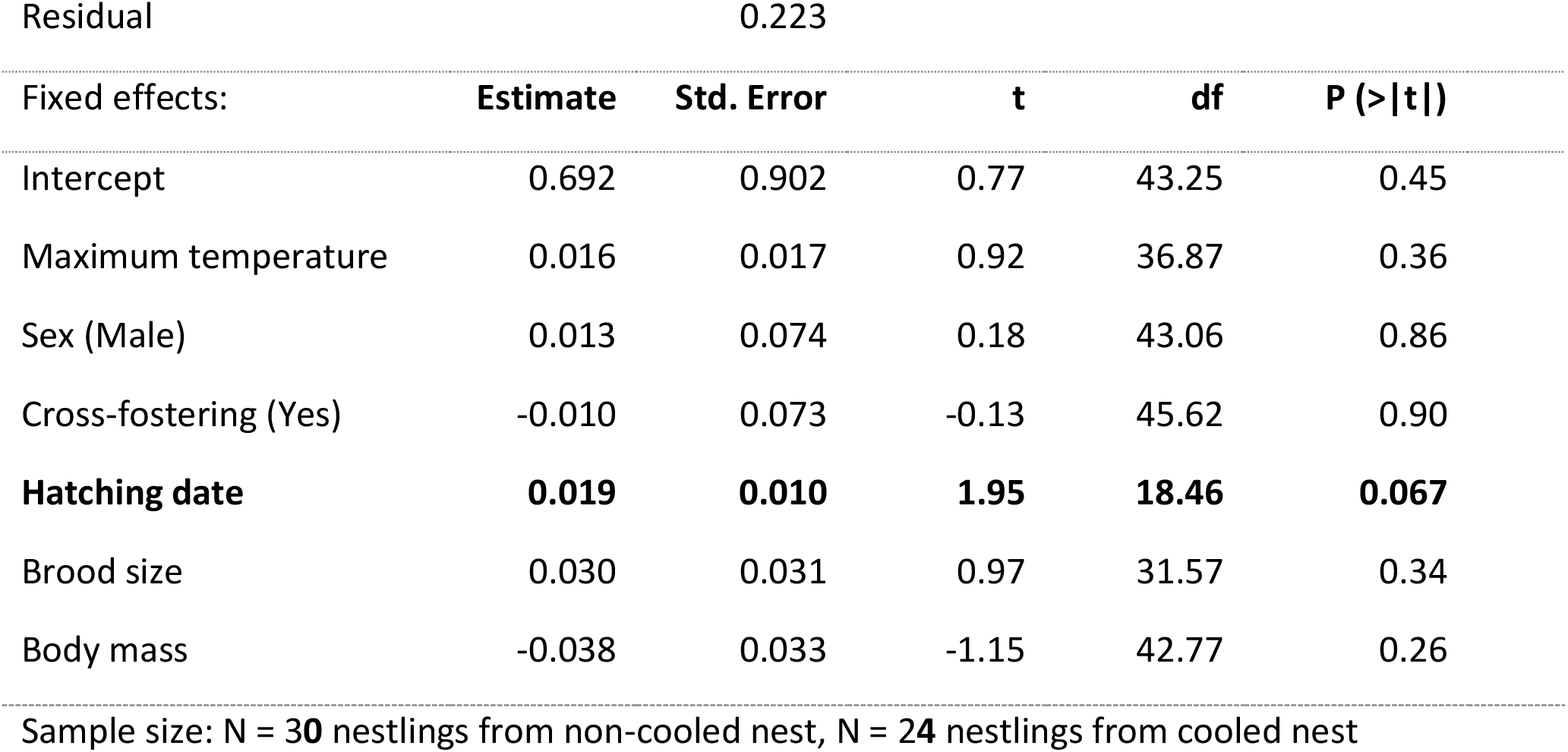
Summary of the linear mixed models testing the relationship between telomere length of nestlings at day 13 after hatching and <u>maximum</u> nestbox temperature. ((A) 2018 heating experiment, (B) 2019 cooling experiment) while controlling for other variables.

