## Supplementary material for "Warm and cool temperatures decrease early-life telomere length in wild pied flycatchers": Table S

**Supplementary Table S1: Summary of the linear mixed models testing the relationship between telomere length of nestlings at day 13 after hatching and mean nestbox temperature** ((A) 2018 heating experiment, (B) 2019 cooling experiment) while controlling for other variables.

| (A) 2018 experimental heating: Day-13 Telomere length (N = 97 nestlings) | | | | | |
|---|---|---|---|---|---|
| Random effects: | | **Variance** | | | |
| Nest of origin (N=26) | Intercept | 0 | | | |
| Nest of rearing (N=41) | Intercept | 0.009 | | | |
| Plate qPCR (N=6) | Intercept | 0.001 | | | |
| Residual | | 0.045 | | | |
| Fixed effects: | **Estimate** | **Std. Error** | **t** | **df** | **P (>\|t\|)** |
| Intercept | 1.395 | 0.451 | 3.09 | 58.33 | 0.003 |
| Mean temperature | -0.012 | 0.015 | -0.80 | 31.32 | 0.43 |
| Sex (Male) | -0.056 | 0.048 | -1.18 | 89.45 | 0.24 |
| Cross-fostering (Yes) | 0.055 | 0.054 | 1.02 | 63.59 | 0.31 |
| Hatching date | 0.003 | 0.004 | 0.67 | 48.81 | 0.50 |
| Brood size | 0.021 | 0.026 | 0.81 | 46.44 | 0.42 |
| Body mass | -0.018 | 0.019 | -0.93 | 78.26 | 0.36 |
| Sample size: N = 55 nestlings from non-heated nest, N = 42 nestlings from heated nest | | | | | |

| (B) 2019 experimental cooling: Day-13 Telomere length (N = 54 nestlings) | | | | | |
|---|---|---|---|---|---|
| Random effects: | | **Variance** | | | |
| Nest of origin (N=2**8**) | Intercept | 0 | | | |
| Nest of rearing (N=4**6**) | Intercept | 0.0002 | | | |
| Plate qPCR (N=10) | Intercept | 0.023 | | | |
| Residual | | 0.053 | | | |
| Fixed effects: | **Estimate** | **Std. Error** | **t** | **df** | **P (>\|t\|)** |
| Intercept | -0.189 | 0.875 | -0.22 | 29.77 | 0.83 |
| **Mean temperature** | **0.068** | **0.032** | **2.12** | **29.29** | **0.043** |
| Sex (Male) | 0.015 | 0.070 | 0.22 | 41.03 | 0.83 |
| Cross-fostering (Yes) | -0.014 | 0.071 | -0.19 | 45.49 | 0.84 |
| **Hatching date** | **0.027** | **0.010** | **2.69** | **16.76** | **0.016** |
| Brood size | -0.001 | 0.032 | -0.04 | 24.80 | 0.97 |

| | | | | | |
|---|---|---|---|---|---|
| Body mass | -0.040 | 0.031 | -1.30 | 40.86 | 0.20 |
| Sample size: N = 30 nestlings from non-cooled nest, N = 24 nestlings from cooled nest | | | | | |

**Supplementary Table S2: Summary of the linear mixed models testing the relationship between telomere length of nestlings at day 13 after hatching and <u>minimum</u> nestbox temperature** ((A) 2018 heating experiment, (B) 2019 cooling experiment) while controlling for other variables.

| (A) 2018 experimental heating: Day-13 Telomere length (N = 97 nestlings) | | | | | |
|---|---|---|---|---|---|
| Random effects: | | **Variance** | | | |
| Nest of origin (N=26) | Intercept | 0 | | | |
| Nest of rearing (N=41) | Intercept | 0.011 | | | |
| Plate qPCR (N=6) | Intercept | 0 | | | |
| Residual | | 0.045 | | | |
| Fixed effects: | **Estimate** | **Std. Error** | **t** | **df** | **P (>\|t\|)** |
| Intercept | 1.139 | 0.385 | 2.96 | 67.69 | 0.004 |
| Minimum temperature | 0.005 | 0.015 | 0.36 | 43.10 | 0.72 |
| Sex (Male) | -0.055 | 0.048 | -1.16 | 89.96 | 0.25 |
| Cross-fostering (Yes) | 0.053 | 0.054 | 0.98 | 68.40 | 0.33 |
| Hatching date | 0.001 | 0.005 | 0.23 | 58.61 | 0.82 |
| Brood size | 0.022 | 0.026 | 0.84 | 49.40 | 0.41 |
| Body mass | -0.019 | 0.019 | -1.00 | 80.70 | 0.32 |
| Sample size: N = 55 nestlings from non-heated nest, N = 42 nestlings from heated nest | | | | | |
| (B) 2019 experimental cooling: Day-13 Telomere length (N = 54 nestlings) | | | | | |
| Random effects: | | **Variance** | | | |
| Nest of origin (N=2**8**) | Intercept | 0 | | | |
| Nest of rearing (N=4**6**) | Intercept | 0.073 | | | |
| Plate qPCR (N=10) | Intercept | 0.129 | | | |
| Residual | | 0.052 | | | |
| Fixed effects: | **Estimate** | **Std. Error** | **t** | **df** | **P (>\|t\|)** |
| Intercept | 0.083 | 0.660 | 1.26 | 30.19 | 0.22 |
| Minimum temperature | 0.046 | 0.032 | 1.41 | 32.75 | 0.17 |

| | | | | | |
|---|---|---|---|---|---|
| Sex (Male) | 0.016 | 0.073 | 0.23 | 42.02 | 0.82 |
| Cross-fostering (Yes) | -0.031 | 0.072 | -0.43 | 45.47 | 0.66 |
| **Hatching date** | **0.019** | **0.009** | **2.17** | **14.82** | **0.047** |
| Brood size | 0.002 | 0.035 | 0.07 | 33.37 | 0.95 |
| Body mass | -0.058 | 0.033 | -1.74 | 42.54 | 0.090 |
| Sample size: N = 30 nestlings from non-cooled nest, N = 24 nestlings from cooled nest | | | | | |

**Supplementary Table S3: Summary of the linear mixed models testing the relationship between telomere length of nestlings at day 13 after hatching and maximum nestbox temperature** ((A) 2018 heating experiment, (B) 2019 cooling experiment) while controlling for other variables.

| (A) 2018 heating experiment: Day-13 Telomere length (N = 97 nestlings) | | | | | |
|---|---|---|---|---|---|
| Random effects: | | **Variance** | | | |
| Nest of origin (N=26) | Intercept | 0 | | | |
| Nest of rearing (N=41) | Intercept | 0.001 | | | |
| Plate qPCR (N=6) | Intercept | 0.001 | | | |
| Residual | | 0.046 | | | |
| Fixed effects: | **Estimate** | **Std. Error** | **t** | **df** | **P (>\|t\|)** |
| Intercept | 1.698 | 0.462 | 3.66 | 58.70 | 0.001 |
| **Maximum temperature** | **-0.017** | **0.009** | **-1.84** | **37.01** | **0.073** |
| Sex (Male) | -0.059 | 0.048 | -1.25 | 89.41 | 0.21 |
| Cross-fostering (Yes) | 0.042 | 0.052 | 0.80 | 64.28 | 0.43 |
| Hatching date | 0.003 | 0.004 | 0.63 | 45.33 | 0.53 |
| Brood size | 0.020 | 0.025 | 0.82 | 43.60 | 0.41 |
| Body mass | -0.020 | 0.019 | -1.09 | 74.13 | 0.29 |
| Sample size: N = 55 nestlings from non-heated nest, N = 42 nestlings from heated nest | | | | | |

| (B) 2019 cooling experiment: Day-13 Telomere length (N = 54 nestlings) | | |
|---|---|---|
| Random effects: | | **Variance** |
| Nest of origin (N=2**8**) | Intercept | 0 |
| Nest of rearing (N=4**6**) | Intercept | 0.103 |
| Plate qPCR (N=10) | Intercept | 0.118 |

| | | | | | |
|---|---|---|---|---|---|
| Residual | | 0.223 | | | |
| Fixed effects: | **Estimate** | **Std. Error** | **t** | **df** | **P (>\|t\|)** |
| Intercept | 0.692 | 0.902 | 0.77 | 43.25 | 0.45 |
| Maximum temperature | 0.016 | 0.017 | 0.92 | 36.87 | 0.36 |
| Sex (Male) | 0.013 | 0.074 | 0.18 | 43.06 | 0.86 |
| Cross-fostering (Yes) | -0.010 | 0.073 | -0.13 | 45.62 | 0.90 |
| **Hatching date** | **0.019** | **0.010** | **1.95** | **18.46** | **0.067** |
| Brood size | 0.030 | 0.031 | 0.97 | 31.57 | 0.34 |
| Body mass | -0.038 | 0.033 | -1.15 | 42.77 | 0.26 |

Sample size: N = 3**0** nestlings from non-cooled nest, N = 2**4** nestlings from cooled nest
